# The fiber knob protein of human adenovirus type 49 mediates highly efficient and promiscuous infection of cancer cell lines using a novel cell entry mechanism

**DOI:** 10.1101/2020.07.20.213223

**Authors:** Alexander T. Baker, Gareth Marlow, James A. Davies, Elise Moses, Rosie M. Mundy, David K. Cole, Pierre J. Rizkallah, Alan L. Parker

## Abstract

The human adenovirus (HAdV) phylogenetic tree is diverse, divided across seven species and comprising over 100 individual types. Species D HAdV are rarely isolated with low rates of pre-existing immunity, making them appealing for therapeutic applications. Several species D vectors have been developed as vaccines against infectious diseases where they induce robust immunity in pre-clinical models and early phase clinical trials. However, many aspects of the basic virology of species D HAdV, including their basic receptor usage and means of cell entry, remain understudied.

Here, we investigated HAdV-D49, which previously has been studied for vaccine and vascular gene transfer applications. We generated a pseudotyped HAdV-C5 presenting the HAdV-D49 fiber knob protein (HAdV-C5/D49K). This pseudotyped vector was efficient at infecting cells devoid of all known HAdV receptors, indicating HAdV-D49 uses an unidentified cellular receptor. Conversely, a pseudotyped vector presenting the fiber knob protein of the closely related HAdV-D30 (HAdV-C5/D30K), differing in four amino acids to HAdV-D49, failed to demonstrate the same tropism. These four amino acid changes resulted in a change in isoelectric point of the knob protein, with HAdV-D49K possessing a basic apical region compared to a more acidic region in HAdV-D30K. Structurally and biologically we demonstrate that HAdV-D49 knob protein is unable to engage CD46, whilst potential interaction with CAR is extremely limited by extension of the DG loop. HAdV-C5/49K efficiently transduced cancer cell lines of pancreatic, breast, lung, oesophageal and ovarian origin, indicating it may have potential for oncolytic virotherapy applications, especially for difficult to transduce tumour types.

**Importance:** Adenoviruses are powerful tools experimentally and clinically. To maximise efficacy, the development of serotypes with low pre-existing levels of immunity in the population is desirable. Consequently, attention has focussed on those derived from species D, which have proven robust vaccine platforms. This widespread usage is despite limited knowledge in their basic biology and cellular tropism.

We investigated the tropism of HAdV-D49, demonstrating it uses a novel cell entry mechanism that bypasses all known HAdV receptors. We demonstrate, biologically, that a pseudotyped HAdV-C5/D49K vector efficiently transduces a wide range of cell lines, including those presenting no known adenovirus receptor. Structural investigation suggests that this broad tropism is the result of a highly basic electrostatic surface potential, since a homologous pseudotyped vector with a more acidic surface potential, HAdV-C5/D30K, does not display a similar pan-tropism. Therefore, HAdV-C5/D49K may form a powerful vector for therapeutic applications capable of infecting difficult to transduce cells.

## Introduction

Human adenoviruses are divided into seven species, A-G(1). These non-enveloped, icosahedral viruses have garnered significant interest as therapeutic vectors since they can be grown and purified to high titers, and because the double-stranded DNA genome is readily amenable to genetic modification, enabling the overexpression of therapeutic transgenes(2, 3). Similar techniques can also be applied to genetically alter the virus structural genes, creating modified viral tropisms which are retained by progeny virions after replication.

Clinically, adenoviruses have been developed as vectors for gene therapy, vaccines, and as oncolytic virotherapies(1, 4, 5). However, efficacy in these applications can be hampered by pre-existing immunity against the therapeutic vector in the population resulting from prior exposure to the wild type pathogen. Such pre-existing immunity is likely to reduce the therapeutic index of such systems, due to rapid and efficient removal of the engineered therapeutic agent by the reticuloendothelial system(6, 7). This is especially relevant where the therapeutic is based on the most commonly studied species C adenovirus, human adenovirus type 5 (HAdV-C5), where neutralising antibodies are found in ~90% of patients from sub-Saharan Africa and ~30% of a Scottish patient cohort(8, 9).

A promising means to circumvent pre-existing immunity is through the development of viruses with naturally low seroprevalence rates as therapeutic agents. For example, vaccines have been developed using chimpanzee adenoviruses which have little to no seroprevalence in humans. However, it appears a significant percentage of some populations may still harbour some immunity to chimpanzee adenoviruses, as observed in a cohort from China(10).

Most attempts to develop adenoviruses with low seroprevalence have focused on those derived from species B or D, due to their comparative rarity(5, 9, 11). The most clinically advanced of these are HAdV-D26 and Enadenotucirev (formerly ColoAd1)(12). Enadenotucirev was developed by evolution of a panel of different adenovirus strains to select for recombinants with rapid replication in tumour cells. The resultant recombinant was predominantly HAdV-B11 with some elements of HAdV-B3, and has progressed into clinical trials as a novel cancer therapeutic(13, 14). HAdV-D26, is a replication deficient vector and the basis of the Ad26.ZEBOV vaccine against Ebola virus, currently under evaluation in the PREVAIL and PREVAC studies(15, 16).

Many species D adenoviruses have previously been evaluated for their potential as vaccines, gene therapies, and oncolytic viruses(1, 5, 9, 11). One with particularly low seroprevalence rates is HAdV-D49. In a cohort of 100 Belgian individuals only 2% had HAdV-D49 positive sera, whilst no pre-existing immunity against HAdV-D49 was detected in 103 Scottish patients(8, 17). Prevalence is somewhat higher in sub-saharan Africa with 22% of 200 patients presenting neutralising antibodies (nAbs), highlighting significant geographical variation in seroprevalence(9).

HAdV-D49 was first isolated from the faeces of a human with no observed disease, and later from Dutch patients(18, 19). It was then isolated from nosocomial epidemic keratoconjunctivitis infections(20, 21), but is most associated with patients who are immunocompromised due to HIV infection(22). A study of adenovirus infections in patients from the UK and Netherlands found 11 instances of HAdV-D49 infection in 183 HIV positive patients (6% HAdV-D49 positive), compared to just two instances in 2301 tested healthy patients (0.09% HAdV-D49 positive)(19).

Previous studies suggest that HAdV-D49 may be effective as a vaccine vector. A vaccine vector based on HAdV-D49 has been evaluated previously for its ability to protect against simian immunodeficiency virus (SIV) challenge. This vector induced strong anti-SIVGag CD8^+^ mediated immunity to SIV, greater than the comparable HAdV-C5 based vector(23). Another study sought to exploit HAdV-D49 a gene therapy to reduce excessive smooth muscle cell proliferation in vascular conduits following bypass grafting. This study demonstrated that HAdV-D49 was efficient at infecting endothelial cells and vascular smooth muscle cells, even after short exposure times(8). Studies in CAR, CD46, and α2-3 linked sialic acid expressing cells have previously suggested that HAdV-D49 may engage CD46 as a cellular receptor, although the effects observed were small(23).

Despite these studies and the development of HAdV-D49 as a therapeutic agent, there remains little information surrounding the basic biology of HAdV-D49 and its means of cellular engagement. Here, we investigate the tropism of HAdV-D49, focussing on the fiber knob protein as the major mediator of cellular attachment, and evaluate the potential utility of a pseudotyped HAdV-C5/D49K vector to infect a range of cancer cell lines.

## Results and Discussion

### HAdV-C5/D49K is not dependent on any known adenovirus receptor for cell entry

To investigate the receptor usage of human adenovirus type 49 fiber knob protein we generated a replication incompetent HAdV-C5 vector pseudotyped with the fiber knob protein of HAdV-D49 (HAdV-C5/D49K), expressing either green fluorescent protein (GFP) or luciferase as transgenes. We also produced a replication deficient HAdV-C5 based pseudotyped vector with the whole fiber protein, including both the fiber shaft and fiber knob of HAdV-D49, expressing luciferase (HAdV-C5/D49F). This pseudotyping approach is a well-established means to investigate the fiber knob in the context of a well understood, replication incompetent virus(1, 24). Using these pseudotyped vectors, we performed transduction assays in CHO cells expressing common adenovirus receptors (Figure 1). CHO-K1 cells do not express any known adenovirus receptor, whilst CHO-CAR cells express the HAdV-C5 receptor, coxsackie and adenovirus receptor (CAR), and CHO-BC1 cells express the BC1 isoform of CD46, the major receptor for species BI adenovirus, which includes HAdV-B35.

**Figure 1:**
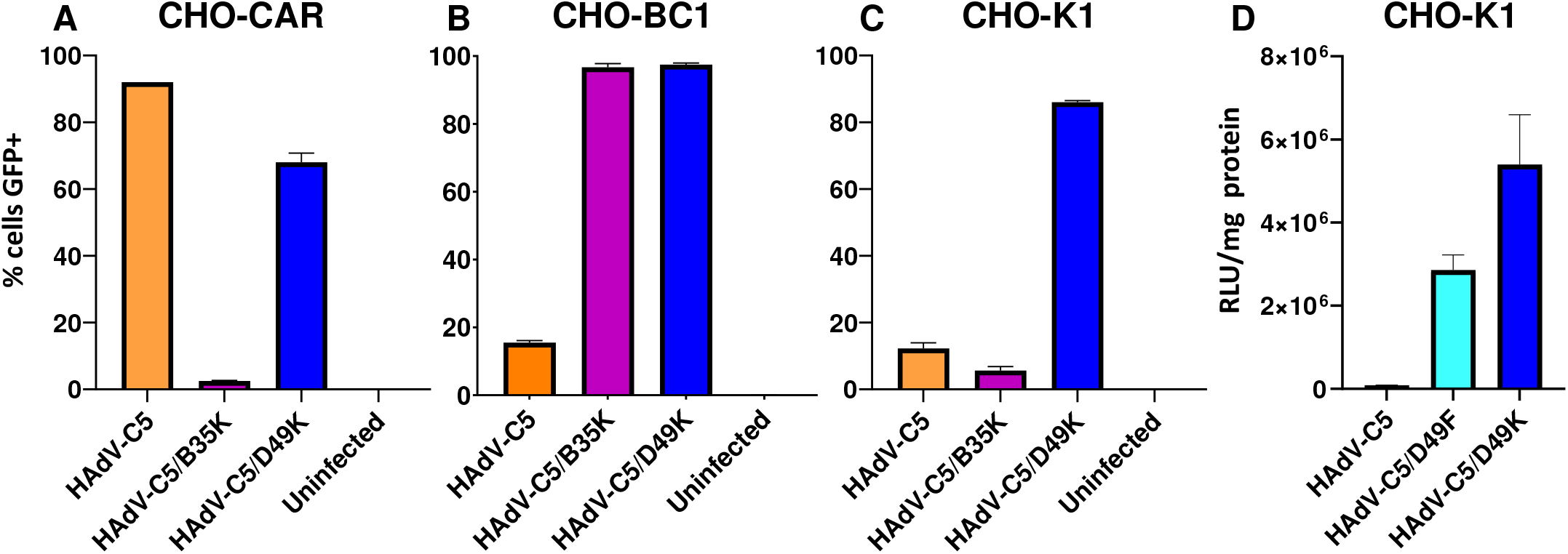
HAdV-C5/D49K infection is not dependent upon CAR or CD46. Transduction assays were performed in Chinese Hamster Ovary (CHO) cells stably expressing CAR (A), CHO-BC1 cells, stably expressing human CD46 isoform BC1 (B), or CHO-K1 cells, which do not express any known adenovirus fiber-knob receptors (C, D). Cells were infected with 5,000 viral particles per cell of replication deficient HAdV-C5, HAdV-C5/B35K, or HAdV-C5/49K expressing a GFP transgene (A-C), or HAdV-C5, HAdV-C5/49K or HAdV-C5/49F expressing luciferase (D). n=3 error is ±SD

The ability of HAdV-C5/D49K to transduce cell lines was compared to similar GFP expressing replication incompetent vectors HAdV-C5 and HAdV-C5/B35K, which engage CAR and CD46 as receptors, respectively(24–26). HAdV-C5/D35K was unable to transduce CHO-CAR, due to the lack of CD46, whilst HAdV-C5 transduced CHO-CAR cells efficiently due to the high levels of CAR expressed. HAdV-C5/D49K transduced CHO-CAR cells efficiently, but slightly less so (by ~20%) compared to HAdV-C5 (Figure 1A). In CHO-BC1 cells, transduction by HAdV-C5 was inefficient, due to the absence of CAR, whilst HAdV-C5/B35 transduced these cells with almost 100% efficiency, due to the presence of high affinity HAdV-B35 receptor, CD46. HAdV-C5/D49K demonstrated a similar ability to HAdV-C5-B35K to transduce CHO-BC1 cells (Figure 1B). In CHO-K1 cells neither HAdV-C5 nor HAdV-C5/B35K were able to efficiently transduce the cells due to the absence of known adenovirus cell surface receptors. HAdV-C5/D49K, however, was able to transduce these cells efficiently, indicating that HAdV-D49K is able to infect cells efficiently and independently of CAR or CD46 (Figure 1C), thus indicating HAdVC5/D49K engages an alternative cellular receptor. Interestingly, we also observed, in this and later experiments, that HAdV-C5/D49K was less efficient at infecting CHO-CAR cells (Figure 1A) compared to non-CAR expressing CHO cell types (Figure 1B-C). These data indicate that the presence of CAR may actively reduce the efficiency of transduction of HAdVC5/D49K compared to levels of transduction in the absence of CAR in the same cell line background.

At 385 amino acids in length, the native HAdV-D49 protein is significantly shorter than the equivalent HAdV-C5 fiber protein, which is 581 amino acids in length. This manifests as a naturally shorter and less flexible fiber shaft in HAdV-D49 compared to that of HAdV-C5. This shortened fiber shaft length can impact upon viral infectivity, resulting in trapping of adenoviral particles within late endosomes due to the decreased endosomolytic activity of shorter shafted adenoviral particles(27). To assess the impact of pseudotyping the entire short fiber protein from HAdV-D49 on viral infectivity, we performed similar transduction assays using CHO-K1 cells. Consistent with engaging an alternative receptor on CHO-K1 cells, the HAdV-C5/D49F whole-fiber pseudotyped vector efficiently transduced CHO-K1 cells, where HAdV-C5 was unable. Also consistent with shorter shafted HAdVs displaying reduced infectivity due to less efficient intracellular trafficking post-entry, the HAdV-C5/D49F was less efficient than the “knob-only” pseudotype HAdV-C5/D49K at infecting CHO-K1 cells (Figure 1D).

We performed similar transduction assays using CHO-K1 and SKOV-3 ovarian cancer cells with and without pre-treatment with either heparinase or neuraminidase to determine the ability of HAdVC5/D49K to bind heparan sulphate proteoglycans (HSPGs) or sialic acid, respectively, to mediate cellular infection (Figure 2). As a positive control for heparinase activity we compared HAdV-C5/D49K infectivity to that of HAdV-C5 in the presence and absence of coagulation factor X (FX), a blood coagulation factor which can facilitate infection of some adenovirus by binding to the viral hexon and cellular heparan sulfate proteoglycans (HSPGs)(28). We observed in CHO-K1 and SKOV-3 cells that transduction levels of HAdV-C5 alone were poor (Figure 2A, B), but were significantly enhanced by the presence of FX, enabling cell entry through cellular HSPGs (29–31). Treatment with heparinase to cleave HSPGs reduced transduction efficiency to that of HAdV-C5 alone. HAdV-C5/D49K transduction efficiency was unaffected by treatment with heparinase (Figure 2A, B), indicating that HAdV-D49 is unlikely to utilise HSPGs for cell entry.

**Figure 2:**
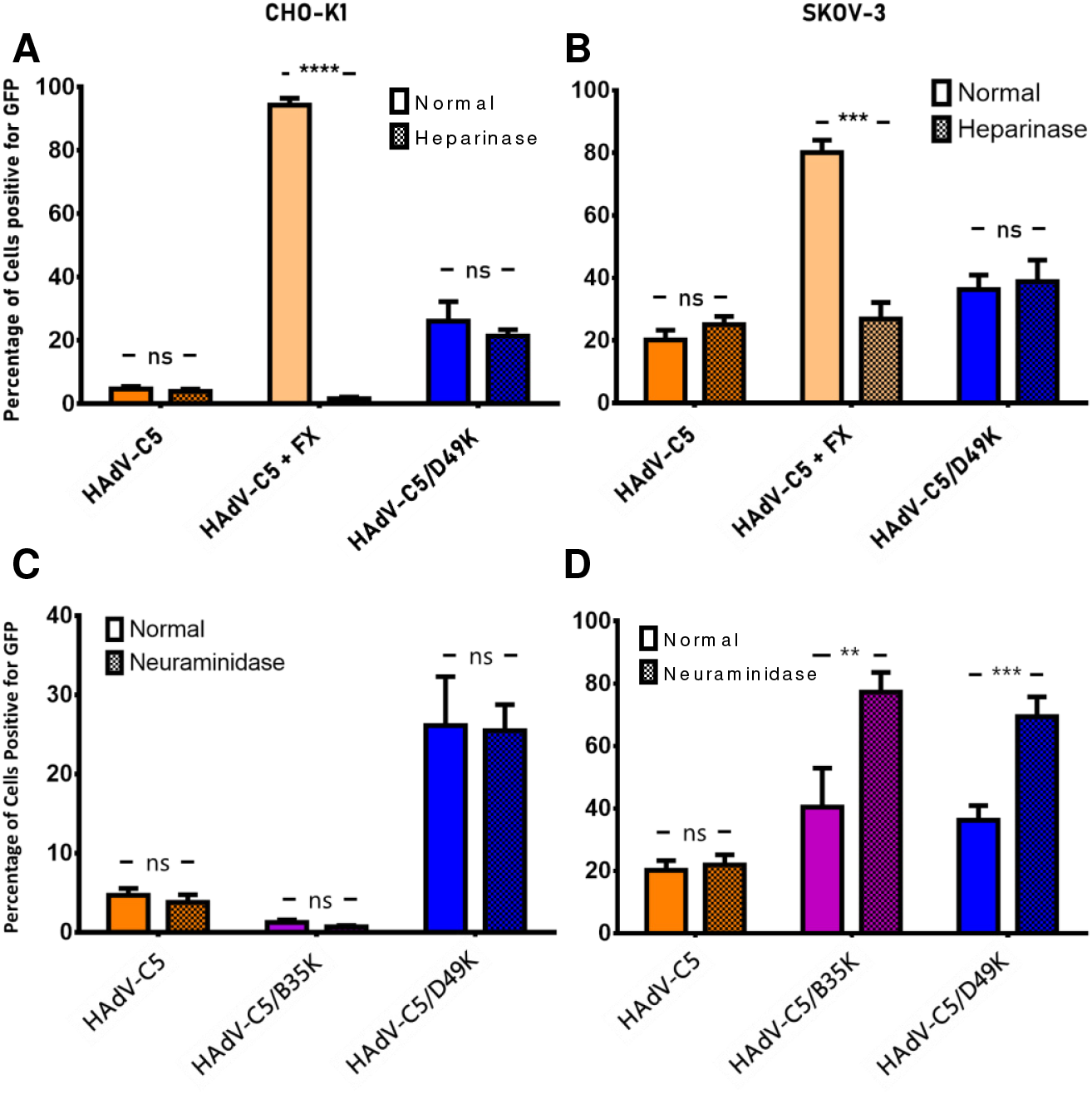
HAdV-C5/49K transduction is not dependent upon HSPGs, sialic acid bearing glycans or Desmoglein 2 (DSG2). Transduction assays were performed in CHO-K1 (A) or SKOV-3 (B) cells with and without heparinase pre-treatment. As a positive control, HAdV-C5 assays also were performed also in the presence of 10μg/ml of FX. Transduction assays were performed with the indicated viral vectors in CHO-K1 (C) or SKOV-3 (D) cells which had been pre-treated with neuraminidase to cleave cell surface sialic acid. Cells were infected with 5,000 viral particles per cell of replication deficient HAdV-C5, HAdV-C5/B35K, or HAdV-C5/49K expressing a GFP transgene, n=3 error is ±SD. *= P<0.05, **= P<0.01, ***= P<0.005,, ****= P<0.001.

Treatment with neuraminidase to remove cellular sialic acid did not alter the ability of any of the viruses to transduce CHO-K1 cells (Figure 2C), as we previously demonstrated to show the involvement of sialic acid in HAdV-C5/D26K infection(24). In SKOV-3 cells, removal of sialic acid actually enhanced the transduction mediated by HAdV-C5/D49K and HAdV-C5/B35K, an effect which we have previously observed by neuraminidase treatment in SKOV-3 cells (Figure 2D)(24, 32, 33). This effect could be a result of the removal of sialic acid enhancing non-specific charge-based interactions between the cell surface and viral capsid. Regardless, these data do not support a role for sialic acid in HAdV-C5/D49K cell infection.

The transduction affinity of HAdV-C5/D49K in the experiments in Figure 2 was noticeably weaker than in the CHO cell experiments (Figure 1). This is due to the methodology used in each experiment. In the transduction experiments (Figure 1), cells were incubated with virus at 37°C for three hours. For studies evaluating the role of sialic acid and HSPGs, cells were pre-treated with enzyme for one hour at 37°C. Then virus was then incubated with cells on ice for one hour following enzymatic digestion to prevent repair and reconstitution of the cleaved heparin/sialic acid. This incubation on ice (and for a shorter period of time) likely decreases viral internalisation during the absorption step, seemingly more profoundly for HAdV-C5/D49K than for the HAdV-C5 suggesting weaker binding at the cell surface or a comparatively low frequency of cell surface receptor.

Desmoglein 2 (DSG2) is the other remaining well-established adenovirus receptor. DSG-2 is described to interact with species BII adenovirus, including HAdV-B3K, via low affinity, avidity dependent mechanism(34). We investigated whether HAdV-D49K might also interact with DSG2 by utilising surface plasmon resonance (SPR), which we have previously used to establish a 66.9μM affinity between DSG2 and HAdV-D3K(35). HAdV-D49K had no detectable affinity for DSG2 (Figure 3A), an unsurprising finding as DSG2 has never been observed as a receptor for any adenovirus outside of species B.

**Figure 3:**
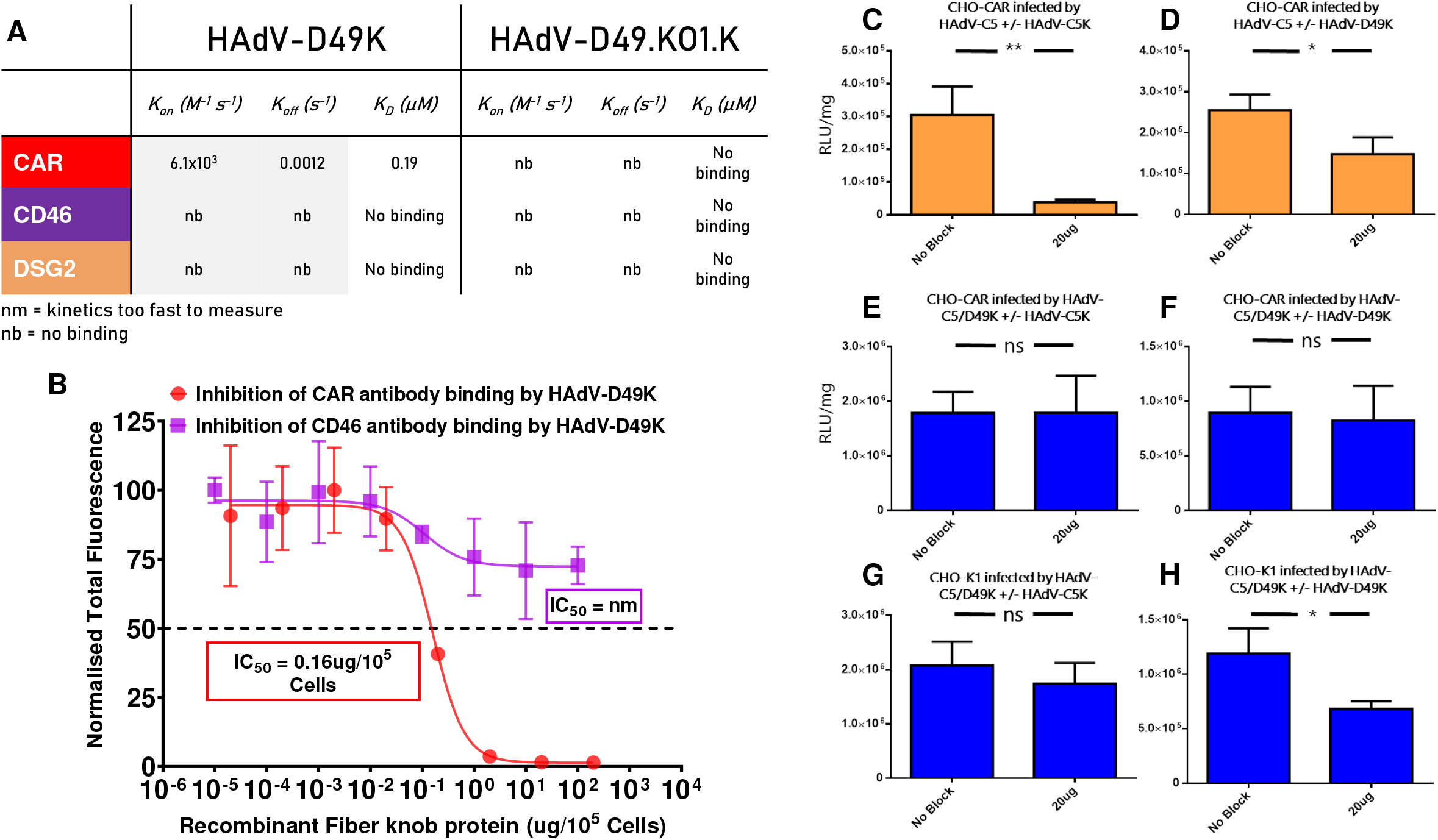
HAdV-D49K interacts with CAR but is not dependent upon it as an entry receptor. Surface plasmon resonance to detect potential interactions between HAdV-D49K or HAdV-D49.KO1.K and CAR, CD46 and DSG2 (A). Antibody binding inhibition assays to assess the ability of recombinant HAdV-D49K to inhibit antibody binding to CHO-CAR or CHO-BC1 cells (B). Blocking of HAdV-C5 mediated transduction was studied by preincubation of CHO-CAR cells with HAdV-C5K (C) or HAdV-D49K (D). Blocking of HAdV-C5/D49K mediated transduction was studied by preincubation of CHO-CAR cells with HAdV-C5K (E) or HAdV-D49K (F). Blocking of HAdV-C5/D49K mediated transduction was studied by preincubation of CHO-K1 cells with HAdV-C5K (G) or HAdV-D49K (H). Cells were infected with 5,000 viral particles per cell of replication deficient HAdV-C5 or HAdV-C5/D49K expressing a luciferase transgene, with and without blockade by 20μg of recombinant HAdV-C5 or HAdV-D49 fiber knob protein. n=3 error is ±SD. *= P<0.05, **= P<0.01.

### HAdV-D49 fiber knob can interact with CAR, but does not require it for cell entry

We also used SPR to further probe the binding affinity of HAdV-D49K, and a mutant version, HAdV-D49.KO1.K, for CD46 and CAR. This HAdV-D49.KO1.K mutant, harbours the KO1 mutations S408E and P409A in the fiber knob AB loop, previously shown to ablate CAR binding in HAdV-C5K(36). The structure of the HAdV-D49.KO1.K fiber knob is also presented (Table 1, PDB 6QPO)(37). As predicted, we did not observe binding between either fiber knob protein and CD46. However, we did observe HAdV-D49K binding to CAR with a detectable 0.19μM affinity which was ablated by the KO1 mutation (Figure 3A).

**Table 1:**
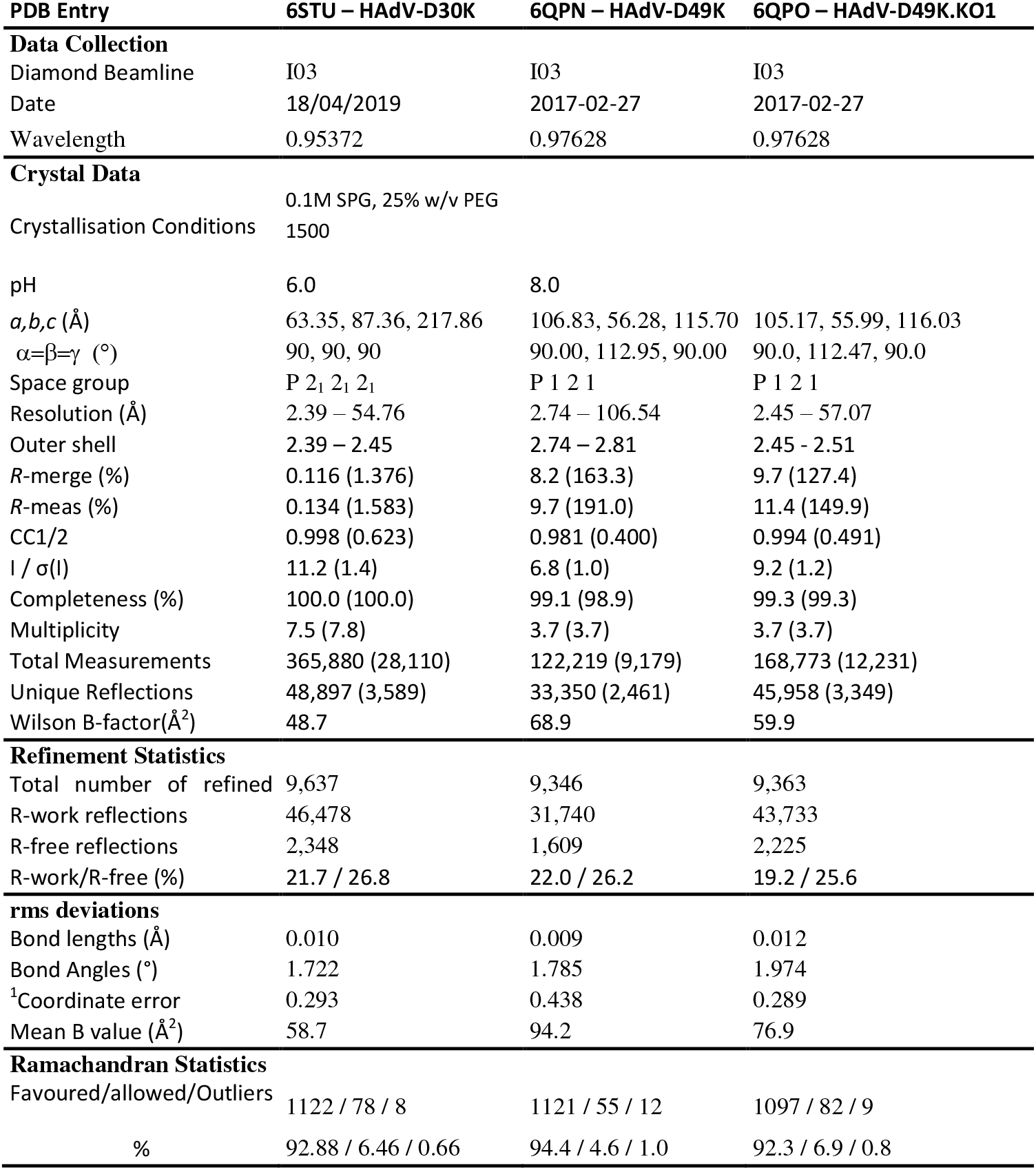
Single crystal diffraction data collection statistics for fiber knob crystal structures determined in this study.

| PDB Entry | 6STU – HAdV-D30K | 6QPN – HAdV-D49K | 6QPO – HAdV-D49K.KO1 |
| --- | --- | --- | --- |
| **Data Collection** |  |  |  |
| Diamond Beamline | I03 | I03 | I03 |
| Date | 18/04/2019 | 2017-02-27 | 2017-02-27 |
| Wavelength | 0.95372 | 0.97628 | 0.97628 |
| **Crystal Data** |  |  |  |
| Crystallisation Conditions | 0.1M SPG, 25% w/v PEG<br>1500 |  |  |
| pH | 6.0 | 8.0 |  |
| *a,b,c* (Å) | 63.35, 87.36, 217.86 | 106.83, 56.28, 115.70 | 105.17, 55.99, 116.03 |
| $\alpha=\beta=\gamma$ (°) | 90, 90, 90 | 90.00, 112.95, 90.00 | 90.0, 112.47, 90.0 |
| Space group | P 2<sub>1</sub> 2<sub>1</sub> 2<sub>1</sub> | P 1 2 1 | P 1 2 1 |
| Resolution (Å) | 2.39 – 54.76 | 2.74 – 106.54 | 2.45 – 57.07 |
| Outer shell | 2.39 – 2.45 | 2.74 – 2.81 | 2.45 - 2.51 |
| *R*-merge (%) | 0.116 (1.376) | 8.2 (163.3) | 9.7 (127.4) |
| *R*-meas (%) | 0.134 (1.583) | 9.7 (191.0) | 11.4 (149.9) |
| CC1/2 | 0.998 (0.623) | 0.981 (0.400) | 0.994 (0.491) |
| *I* / $\sigma(I)$ | 11.2 (1.4) | 6.8 (1.0) | 9.2 (1.2) |
| Completeness (%) | 100.0 (100.0) | 99.1 (98.9) | 99.3 (99.3) |
| Multiplicity | 7.5 (7.8) | 3.7 (3.7) | 3.7 (3.7) |
| Total Measurements | 365,880 (28,110) | 122,219 (9,179) | 168,773 (12,231) |
| Unique Reflections | 48,897 (3,589) | 33,350 (2,461) | 45,958 (3,349) |
| Wilson B-factor(Å<sup>2</sup>) | 48.7 | 68.9 | 59.9 |
| **Refinement Statistics** |  |  |  |
| Total number of refined | 9,637 | 9,346 | 9,363 |
| R-work reflections | 46,478 | 31,740 | 43,733 |
| R-free reflections | 2,348 | 1,609 | 2,225 |
| R-work/R-free (%) | 21.7 / 26.8 | 22.0 / 26.2 | 19.2 / 25.6 |
| **rms deviations** |  |  |  |
| Bond lengths (Å) | 0.010 | 0.009 | 0.012 |
| Bond Angles (°) | 1.722 | 1.785 | 1.974 |
| <sup>1</sup>Coordinate error | 0.293 | 0.438 | 0.289 |
| Mean B value (Å<sup>2</sup>) | 58.7 | 94.2 | 76.9 |
| **Ramachandran Statistics** |  |  |  |
| Favoured/allowed/Outliers | 1122 / 78 / 8 | 1121 / 55 / 12 | 1097 / 82 / 9 |
| % | 92.88 / 6.46 / 0.66 | 94.4 / 4.6 / 1.0 | 92.3 / 6.9 / 0.8 |

We performed IC_50_ binding studies using recombinant HAdV-D49K protein on CHO-CAR and CHO-BC1 cells and assessed the ability of the recombinant fiber knob protein to inhibit the binding of anti-CAR or CD46 antibodies, respectively (Figure 3B). HAdV-D49K was able to block anti-CAR antibody binding to CHO-CAR cells in a dose dependent manner (IC_50_ = 0.16μg/10^5^ cells, Figure 3B). However, no IC_50_ could be derived by using HAdV-D49K to block CD46 on CHO-BC1 cells, where HAdV-D49K was unable to achieve more than 20% inhibition of antibody binding, suggesting weak or incidental CD46 interactions (Figure 3B). Therefore, these data support the findings from SPR and transduction experiments that HAdV-D49K may bind CAR with low affinity, but does not bind CD46.

Our earlier findings indicated that HAdV-C5/D49K is not dependent upon CAR for cell entry (Figure 1), however our *in vitro* biological inhibition and SPR assays demonstrate CAR binding affinity (Figure 3A, B).

We further investigated this finding by performing transduction blocking experiments using the CAR engaging HAdV-C5 and HAdV-C5/D49K with recombinant fiber knob protein of each virus in CHO-CAR and CHO-K1 cells. As predicted, pre-incubation of CHO-CAR cells with recombinant HAdV-C5K efficiently inhibited HAdV-C5 infection (Figure 3C), whilst blocking with HAdV-D49K inhibited infection by HAdV-C5 by approximately 50% (Figure 3D). Infecting CHO-CAR cells with HAdV-C5/D49K pseudotype and attempting to block using HAdV-C5K (Figure 3E) or HAdV-D49K (Figure 3F) did not significantly inhibit transduction efficiency. Finally, infection of CHO-K1 cells by HAdV-C5/D49K and blocking with HAdV-C5K did not significantly inhibit transduction efficiency (Figure 3G), whilst blocking with HAdV-D49K reduced transduction efficiency by approximately 50% (Figure 3H).

These data confirm that HAdV-D49K is capable of binding to CAR, albeit at approximately 1000x lower affinity than HAdV-C5(35), but in a manner able to inhibit HAdV-C5 binding. These data confirm HAdV-C5/D49K is capable of entering cells though a non-CAR mediated pathway, since HAdV-C5K cannot inhibit HAdV-C5/D49K transduction in CHO-CAR cells (Figure 3E). Interestingly, HAdV-C5/D49K was able to inhibit its own viral infection only in the absence of CAR (Figure 3H).

One potential explanation for this activity is that the unknown alternative receptor to CAR has a lower affinity for HAdV-D49K than CAR. Therefore, in the presence of CAR the recombinant fiber knob would be sequestered on the higher affinity CAR receptor leaving the alternative receptor free to interact with the virus. A low affinity receptor would also, likely, depend upon avidity, so might not be observed with single trimers of HAdV-D49K; a similar effect has previously been observed with HAdV-B3K and DSG2(38) and CD46(39). This is supported by the observation what HAdV-D49K cannot transduce cells as efficiently when incubated on ice in the absence of CAR, while HAdV-C5 and HAdV-C5/B35K, which form high affinity receptor interactions, are unencumbered (Figure 2).

### HAdV-D49K may bind cells through a charge dependent mechanism

To investigate other closely related HAdV with homologous fiber-knob proteins we performed a BLASTp search using the HAdV-D49K amino acid sequence. This search revealed that the HAdV-D30K protein is highly homologous to HAdV-D49K, differing in just 4 amino acid residues (Figure 4A).

**Figure 4:**
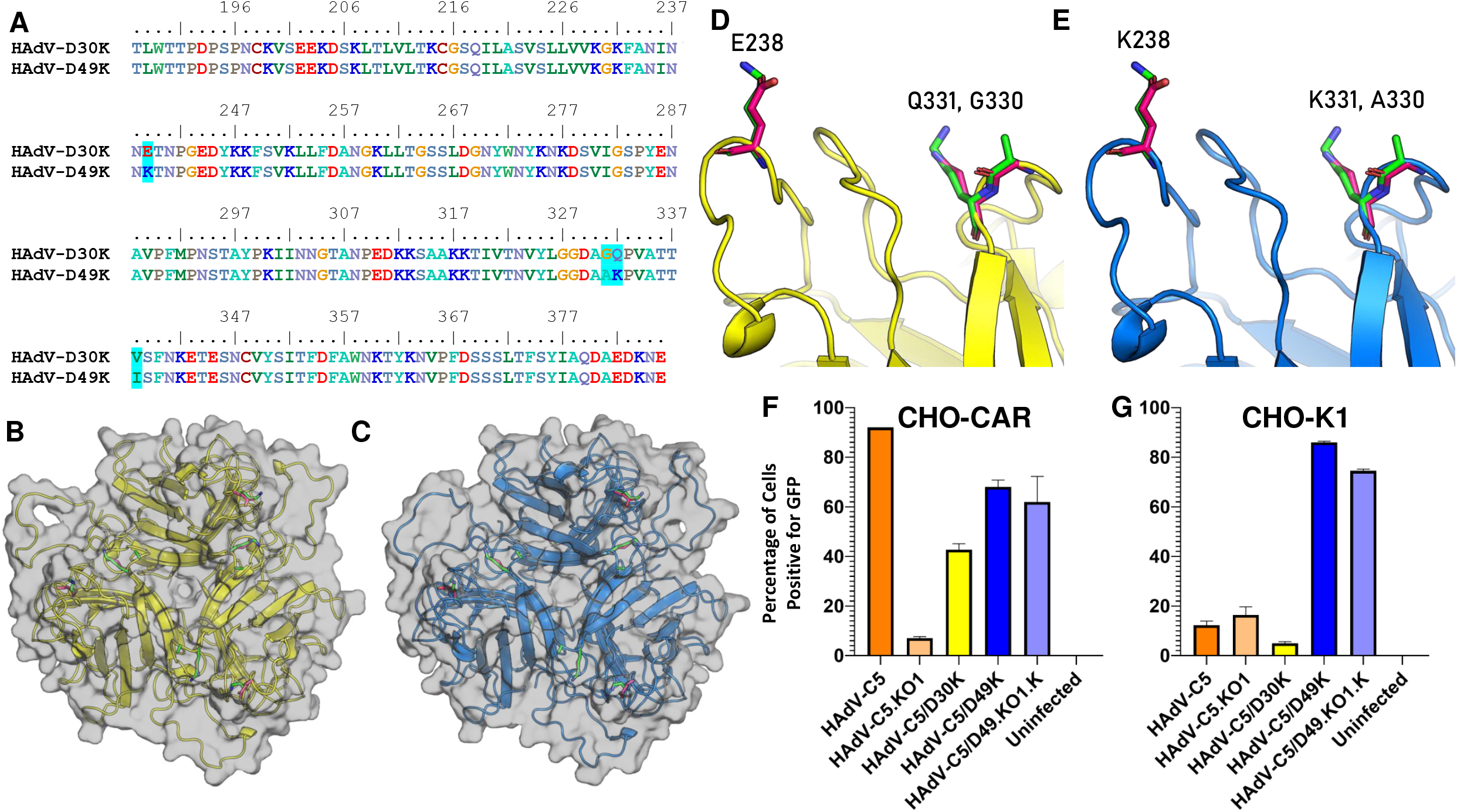
HAdV-D49K differs from HAdV-D30K in only three surface exposed amino acids but demonstrates radically altered cellular tropism. Clustal Ω sequence alignment (numbering based on whole fiber sequence) of HAdV-D30K and HAdV-D49K (A). Viewed from the apex down the three fold axis, as if towards the viral capsid, the crystal structures of HAdV-D30K (B) and HAdV-D49K (C) reveal that three of these residues are surface exposed. These residues can be seen projecting into the solvent from loops on the apex of HAdV-D30K (D) and HAdV-D49K (E), residue numbers and names correspond to the fiber-knob protein depicted in that frame. Sticks representing residues belonging to HAdV-D30K and HAdV-D49K are seen in pink and green, respectively. Transduction assays were performed to assess tropism of HAdV-C5/D30K, HAdV-C5/D49K and HAdV-C5/D49.KO1.K in CHO-CAR cells (F) and CHO-K1 cells (G). Cells were infected with 5,000 viral particles per cell of replication deficient HAdV-C5 or HAdVC5/D49K expressing a luciferase transgene, with and without blockade by 20μg of recombinant HAdV-C5 or HAdV-D49 fiber knob protein.

We solved the crystal structures of HAdV-D30K (Figure 4B) and HAdV-D49K (Figure 4C). Diffraction data collection statistics for these structures are provided in Table 1. We demonstrate that structurally, HAdV-D30K and HAdV-D49K are highly homologous (RMSD = 0.292Å^2^). Residue 338 is not surface exposed on either fiber knob protein and is likely to be functionally homologous (HAdV-D49 = Isoleucine338, HAdV-D30 = Valine338). However, the remaining 3 residue differences, E339K, G331A, and Q332K (HAdV-D30K → D49K) are surface exposed at the apex of each fiber knob monomer (Figure 4D, E). The E339K and Q332K substitutions have opposing charges.

We investigated the transduction efficiency of HAdV-C5/D30K compared to HAdV-C5/D49K, HAdV-C5, HAdV-C5.KO1, and HAdV-C5/D49.KO1.K in CHO-CAR (Figure 4F) and CHO-K1 cells (Figure 4G). HAdV-C5 infected CHO-CAR cells efficiently whilst the CAR-binding ablated KO1 mutant version did not, whilst HAdV-C5/49K and the corresponding mutant HAdV-C5/49KO1.K infected CHO-CAR cells with similar efficiency as observed for HAdV-C5/D49K in Figure 1A. HAdV-C5/D30K infected CHO-CAR cells with approximately 40% efficiency (Figure 4F). In CHO-K1 cells HAdV-C5/D49K and the KO1 mutant were the only viruses which achieved efficient transduction. Surprisingly, given the high homology to HAdVC5/D49K, the HAdV-C5/D30K pseudotype was inefficient in transducing CHO-K1 cells (<5% GFP^+^, Figure 4G).

This profound difference in transduction efficiency between HAdV-C5/D30K and HAdV-C5/D49K must be dependent upon the 3 surface exposed amino acid differences. We investigated the effect of the opposing charges at residue substitutions 339 and 332 (Figure 4A-E) by modelling the electrostatic surface potential of the two fiber knob proteins, based on our crystal structures (Figure 5). The surface potential maps reveal that whilst structural homology was high, they present radically different electrostatic surface potential distributions. HAdV-D30K is significantly more acidic (pI = 5.57) than HAdV-D49K (pI = 8.26) (Figure 5).

**Figure 5:**
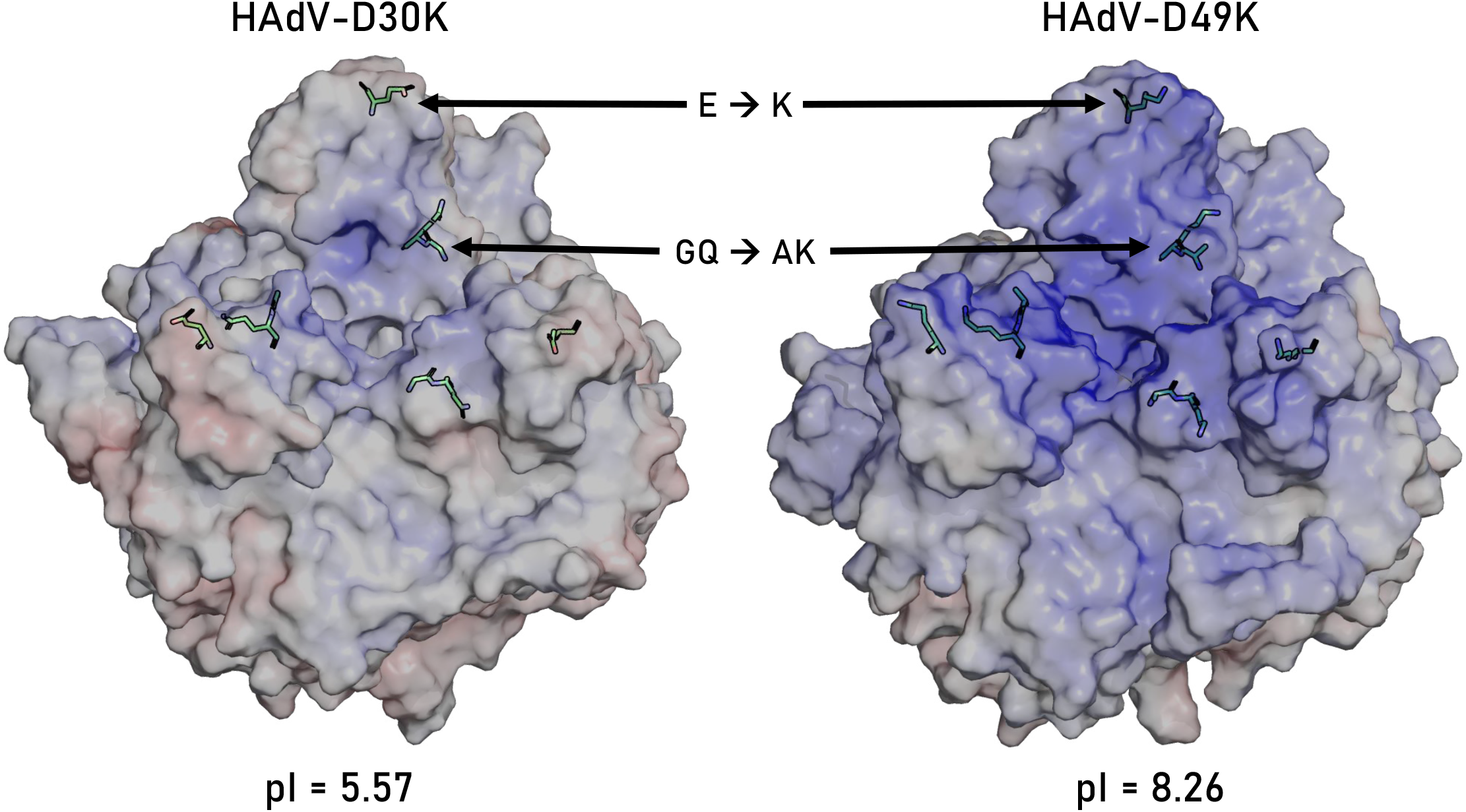
The residue differences between HAdV-D30K and HAdV-D49K effect the surface electrostatic potential of the fiber knob. The calculated pI of HAdV-D30K and HAdV-D49K is differ as a result of the residue changes, which are shown as green sticks as they occur in that fiber knob protein. The calculated electrostatic surface potential at pH7.35 is projected on a −10mV to +10mV ramp (Red to Blue). HAdV-D49K is seen to have much more basic potential around the apex where the residue substitutions are.

Thus, it seems probable that the interaction with the unknown cell surface receptor requires basic electrostatic potential. This is commensurate with the previous inference that its receptor is likely to be low affinity, as electrostatic interfaces are often observed to be less stable than their ionic counterparts. It is possible that the electrostatic potential differences explain the reduced transduction affinity observed in HAdV-C5/D30K compared to HAdV-C5/D49K in CHO-CAR cells. Should the strong charge on HAdV-D49K be opposed to that on the surface of CAR, this could enhance the interaction stability and therefore overall virus affinity. It seems unlikely that the residue substitutions themselves would strongly influence CAR affinity as they occur at the apex of the fiber knob, an area which is not critical the CAR interface(1).

### HAdV-C5/D49K is able to efficiently infect a large range of cancer cell lines

Given HAdV-C5/D49K infects cells independently of known adenovirus receptors we hypothesised that it may form the basis of an efficient vector for cancer virotherapy applications. We therefore compared its transduction efficiency to that of HAdV-C5 in panels of pancreatic, breast, oesophageal, colorectal, ovarian and lung cancer cell lines (Table 2).

**Table 2:**
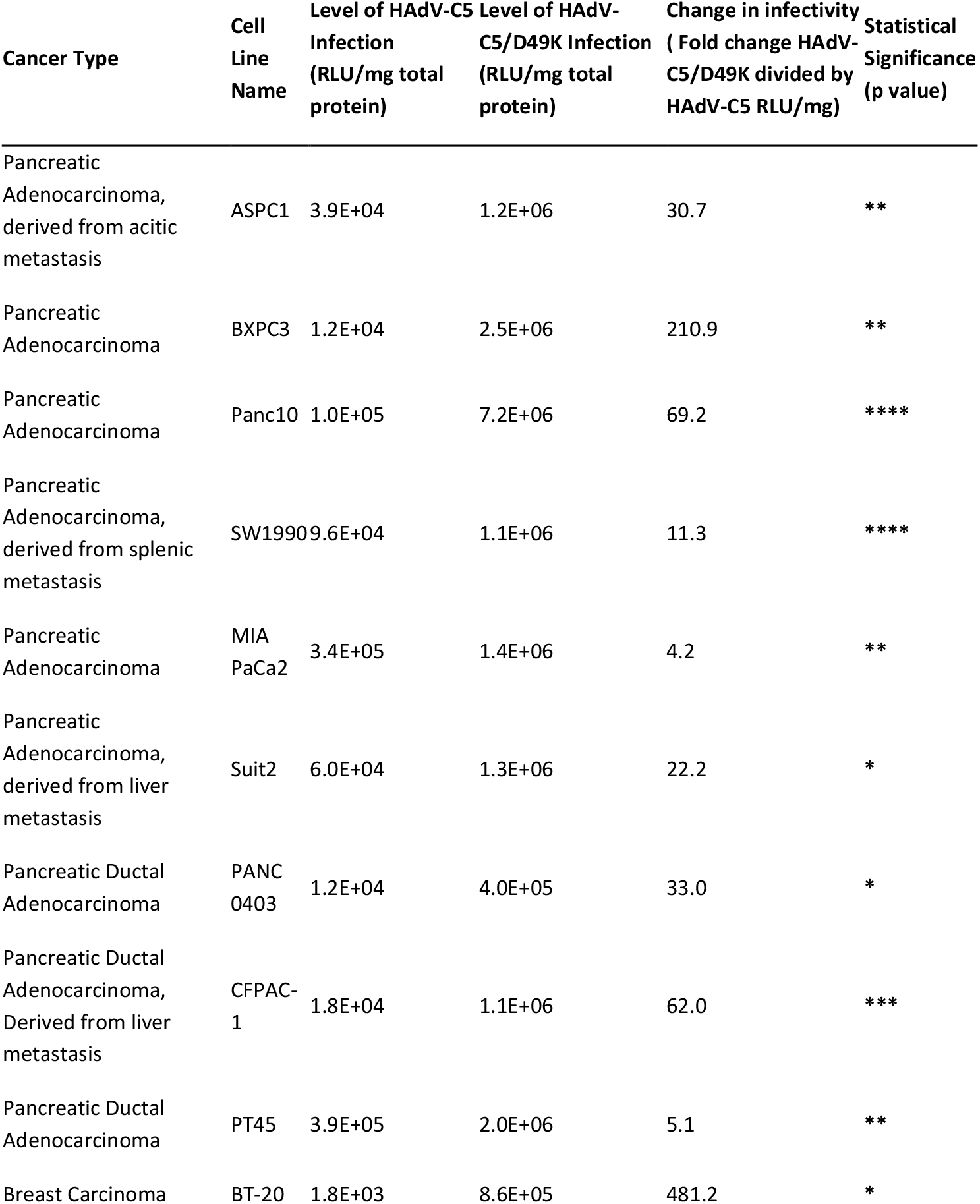
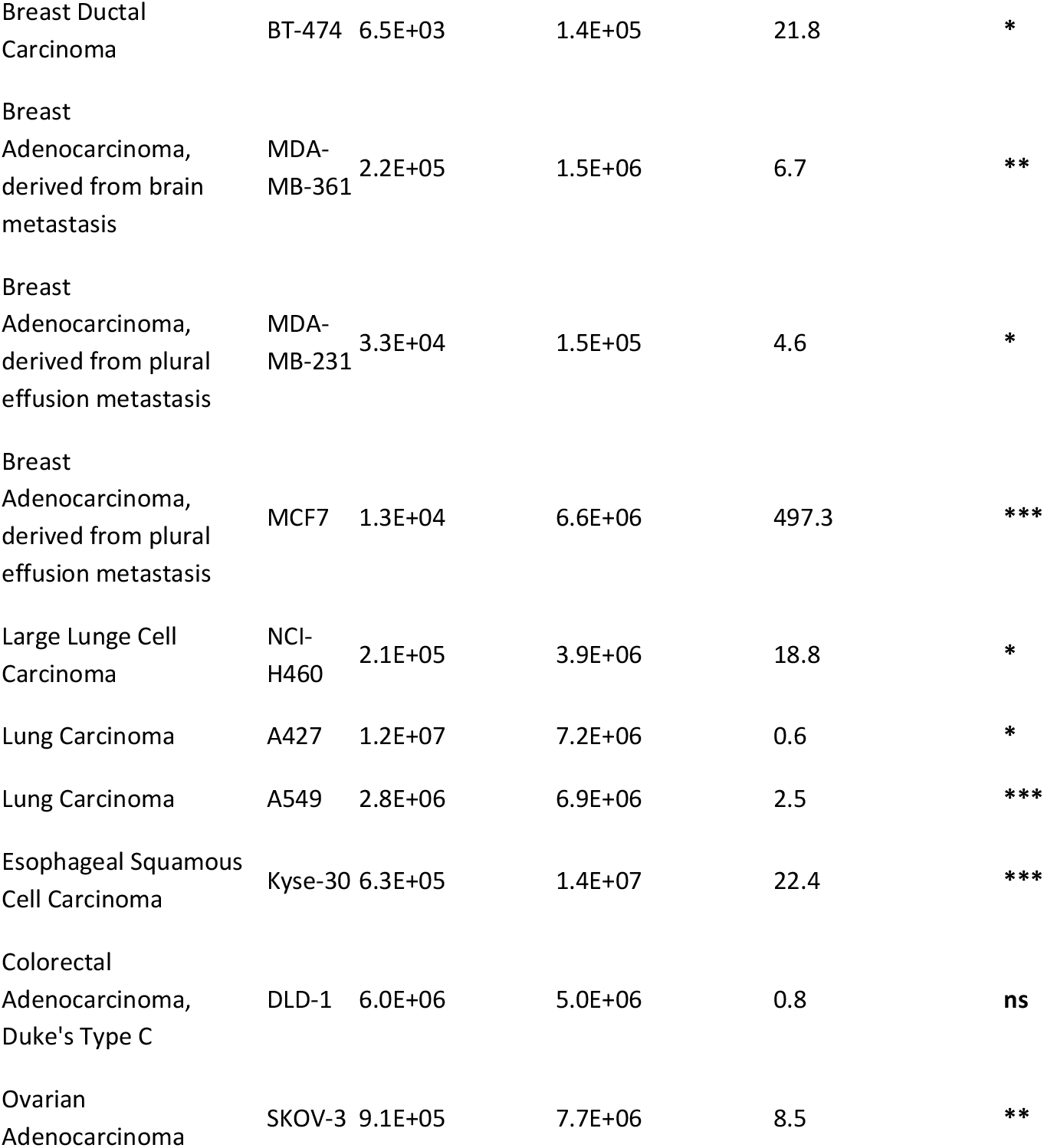
Comparison of transduction of various cancer cell lines by HAdV-C5 and HAdV-C5/D49K. *= P<0.05, **=P<0.01, ***=P<0.001, ****=P<0.0001, ns=not significant.

| Cancer Type | Cell Line Name | Level of HAdV-C5 Infection (RLU/mg total protein) | Level of HAdV-C5/D49K Infection (RLU/mg total protein) | Change in infectivity ( Fold change HAdV-C5/D49K divided by HAdV-C5 RLU/mg) | Statistical Significance (p value) |
| --- | --- | --- | --- | --- | --- |
| Pancreatic Adenocarcinoma, derived from acitic metastasis | ASPC1 | 3.9E+04 | 1.2E+06 | 30.7 | ** |
| Pancreatic Adenocarcinoma | BXPC3 | 1.2E+04 | 2.5E+06 | 210.9 | ** |
| Pancreatic Adenocarcinoma | Panc10 | 1.0E+05 | 7.2E+06 | 69.2 | **** |
| Pancreatic Adenocarcinoma, derived from splenic metastasis | SW1990 | 9.6E+04 | 1.1E+06 | 11.3 | **** |
| Pancreatic Adenocarcinoma | MIA PaCa2 | 3.4E+05 | 1.4E+06 | 4.2 | ** |
| Pancreatic Adenocarcinoma, derived from liver metastasis | Suit2 | 6.0E+04 | 1.3E+06 | 22.2 | * |
| Pancreatic Ductal Adenocarcinoma | PANC 0403 | 1.2E+04 | 4.0E+05 | 33.0 | * |
| Pancreatic Ductal Adenocarcinoma, Derived from liver metastasis | CFPAC-1 | 1.8E+04 | 1.1E+06 | 62.0 | *** |
| Pancreatic Ductal Adenocarcinoma | PT45 | 3.9E+05 | 2.0E+06 | 5.1 | ** |
| Breast Carcinoma | BT-20 | 1.8E+03 | 8.6E+05 | 481.2 | * |
| Breast Ductal Carcinoma | BT-474 | 6.5E+03 | 1.4E+05 | 21.8 | * |
| Breast Adenocarcinoma, derived from brain metastasis | MDA-MB-361 | 2.2E+05 | 1.5E+06 | 6.7 | ** |
| Breast Adenocarcinoma, derived from plural effusion metastasis | MDA-MB-231 | 3.3E+04 | 1.5E+05 | 4.6 | * |
| Breast Adenocarcinoma, derived from plural effusion metastasis | MCF7 | 1.3E+04 | 6.6E+06 | 497.3 | *** |
| Large Lunge Cell Carcinoma | NCI-H460 | 2.1E+05 | 3.9E+06 | 18.8 | * |
| Lung Carcinoma | A427 | 1.2E+07 | 7.2E+06 | 0.6 | * |
| Lung Carcinoma | A549 | 2.8E+06 | 6.9E+06 | 2.5 | *** |
| Esophageal Squamous Cell Carcinoma | Kyse-30 | 6.3E+05 | 1.4E+07 | 22.4 | *** |
| Colorectal Adenocarcinoma, Duke's Type C | DLD-1 | 6.0E+06 | 5.0E+06 | 0.8 | ns |
| Ovarian Adenocarcinoma | SKOV-3 | 9.1E+05 | 7.7E+06 | 8.5 | ** |

In pancreatic cancer cell lines HAdV-C5/D49K was consistently more efficient at cellular transduction than HAdV-C5. This improved activity ranged between 4.2x more efficient in MiaPaCa2 cells to 210.9x more efficient in BxPc3 cells. The most effectively transduced cell line was Panc10 cells, producing 7.2×10^6^ RLU/mg of fluorescence, compared to the least efficient at just 4.0×10^5^ RLU/mg in Panc0403 cells. This suggests that the large differences between HAdV-C5/D49K and HAdV-C5 transduction levels are likely due to the variability in the expression of CAR. A similarly broad range of different relative infection efficiencies were observed in the breast cancer cell lines studied. In MCF7 cells and BT20 cells HAdV-C5/D49K was nearly 500-fold more efficient at transducing the cells due to these cells expressing low levels of CAR, with consequent poor levels of HAdV-C5 mediated transduction (Table 2).

DLD-1 colorectal cancer cells were efficiently transduced by both HAdV-C5 and HAdV-C5/D49K vectors, at 5.0×10^6^ and 6.0×10^6^ RLU/mg respectively. This is the only cell line where no significant difference in infectivity was observed. A427 lung carcinoma cells are the only line where HAdV-C5 mediated cellular transduction was more efficient than HAdV-C5/D49K (0.6x). Whilst HAdV-C5/D49K transduced A427 cell very efficiently (7.2×10^6^ RLU/mg) HAdV-C5 achieved unusually high transduction efficiency (1.2×10^7^ RLU/mg) (Table 2). Therefore, this result is not due to inefficient HAdV-C5/D49K transduction, but unusually efficient HAdV-C5 transduction.

## Conclusions

Previous experiments using whole HAdV-D49 virus concluded that it utilises CD46 as its primary cellular receptor(23). The data described, generated using either purified HAdV-D49 fiber knob protein or a pseudotyped HAdV-C5/D49K vector, clearly demonstrate CD46 to be implausible as a receptor for HAdV-D49.

We demonstrate that the HAdV-D49 fiber knob has a weak affinity for CAR, although it is not dependent upon this interaction to mediate efficient cell entry, and in fact the presence of CAR may inhibit cellular transduction. In support of this we show a mutant vector, HAdV-C5/D49.KO1.K, containing mutations within the fiber knob domain which ablate CAR affinity, retained the ability to efficiently transduce cells in the absence of any detectable binding to CAR. Based on the low efficiency by which HAdV-C5/D49K transduced cells when absorbed on ice and the observation that HAdV-D49K is only capable of inhibiting HAdV-C5/D49K transduction in the absence of CAR, we tentatively suggest that the unknown receptor is likely to be bound with weak affinity and virus attachment may be avidity dependent.

Regardless of the mechanism of interaction this study strongly suggests there is an as yet unknown adenovirus receptor or mechanism of cell entry which mediates efficient transduction of a broad range of cell lines. This is demonstrated by its ability to efficiently infect every cell line tested, throughout this study. The weakest observed transduction was in Panc0403 cells where it achieved 4.0×10^5^ RLU/mg of luminescence. Whilst this is not a particularly strong transduction efficiency, it is still significantly higher (33.0x, P<0.05) than that of HAdV-C5. It is likely, therefore, that the HAdV-C5/D49K vector described here may be useful in biotechnology applications to efficiently express proteins in difficult to transduce cell lines.

HAdV-C5/D49K represents a highly efficient gene transfer vehicle which is not restricted by any known adenovirus tropism. It possesses a broad range of infectivity and has potential as both a laboratory reagent, for the transient expression of transgenes, and as a therapeutic vaccine or oncolytic virus. For oncolytic applications, it is likely that further refinement, such as the introduction of mutations known to confer tumour selective replication, such as dl24 mutation(40–42) or the use of tumour specific promoters such as hTERT(43) or survivin(44) to drive transgene therapeutic expression selectively within tumour cells will be necessary to ensure tight tumour selectivity.

## Materials and Methods

### GFP transduction assay

Adherent cells were seeded into a Nunc delta surface 96-well cell culture plate (ThermoFisher) at a density of 5×10^4^ cells/well in 200μl of cell culture media and left to adhere overnight at 37°C in a 5% CO_2_ humidified atmosphere. Media was removed and cells washed twice with 200μl of PBS. Virus was added at the desired concentration in 200μl of serum free RMPI 1640 and incubated for 3hrs. The virus containing media was then removed and replaced with complete cell culture media and the cells incubated for a further 45hrs. Cell culture media was then removed, the cells washed twice with 200μl of PBS, trypsinised in 50μl of 0.05% Trypsin-EDTA (Gibco), and dissociated by pipetting. The trypsinised cells were transferred to a 96-well V-bottom plate (ThermoFisher), neutralised with 100μl of complete cell culture media, and pelleted by centrifugation at 1200RPM for 3mins. Supernatant was removed, the cells washed once in 200μl of PBS, and resuspended in 100μl of 2% PFA (PBS containing 2% w/v paraformaldehyde) and incubated at 4°C for 15mins. Cells were again pelleted, washed twice in 200μl PBS, then resuspended in 200μl PBS prior to analysis by flowcytometry.

Samples were analysed by flow cytometry on Attune NxT (ThermoFisher), voltages were set prior to each experiment, for each cell type, using an uninfected cell population treated identically. Data was analysed using FlowJo v10 (FlowJo, LLC), gating sequentially on singlets, cell population, and GFP positive cells. Levels of infection were defined as the percentage of GFP positive cells (% +ve), and/or Total Fluorescence (TF), defined as the percentage of GFP positive cells multiplied by the median fluorescent intensity (MFI) of the GFP positive population. These measures are distinct in that % +ve describes the total proportion of cells infected, and TF describes the total efficiency of transgene delivery.

### Luciferase transduction assay

Luciferase infectivity assays were performed using the luciferase assay system kit (Promega). Cells were seeded into a Nunc delta surface 96-well cell culture plate (ThermoFisher) at a density of 2×10^4^ cells/well in 200μl of cell culture media and left to adhere overnight at 37°C in a 5% CO_2_ humidified atmosphere. Media was removed and cells washed once with 200μl of PBS. Luciferase transgene encoding replication incompetent viruses were added to the wells at the required titre in 200μl of serum free RMPI 1640 and incubated for 3hrs. The virus containing media was then removed and replaced with complete cell culture media and the cells incubated for a further 45hrs. Cell culture media was then removed, the cells washed twice with 200μl of PBS, and were then lysed in 100μl of cell culture lysis buffer (part of the Promega kit) diluted to 1x in ddH_2_O. The plate was then frozen at −80°C.

After thaw, 10μl of lysate from the cell culture plate mixed then was transferred to a white Nunc 96-microwell plate (ThermoFisher) and 100μl of luciferase assay reagent (Promega Kit) added. Luciferase activity was then measured in relative light units (RLU) by plate reader (Clariostar, BMG Labtech). Total protein concentration was determined in the lysate using the Pierce BCA protein assay kit (Thermofisher) according to the manufacturers protocol, absorbance was measured on an iMark microplate absorbance reader (BioRad).

Relative virus infection was determined by normalising the measured luciferase intensity to the total protein concentration (RLU was divided by protein concentration). This gave a final infectivity readout in RLU/mg of protein.

### Blocking of virus infection with recombinant fiber knob protein

This assay was also performed using the luciferase assay system kit (Promega). Cells were seeded into a Nunc delta surface 96-well cell culture plate (ThermoFisher) at a density of 2×10^4^ cells/well in 200μl of cell culture media and left to adhere overnight at 37°C in a 5% CO_2_ humidified atmosphere. Media was removed and cells washed 2x with 200μl of cold PBS and the plate cooled on ice. 20pg/cell of recombinant adenovirus fiber knob was added to each well in 200μl of cold PBS and incubated on ice in a 4°C cold room for 1hr. Media was then removed and luciferase transgene encoding replication incompetent viruses added to the necessary wells at the required titre in 200μl of cold serum free RMPI 1640 and incubated on ice in a 4°C cold room for 1hr. The virus containing media was then removed and replaced with complete cell culture media and the cells incubated for a further 45hrs under normal cell culture conditions. From this point forward the assay is identical to the GFP and luciferase transduction assays.

### Heparinase and Neuraminidase transduction assays

Cells were seeded at a density of 5×10^4^ cells/well in a flat bottomed 96 well cell culture plate and incubated overnight at 37°C to adhere. Cells were washed twice with 200μl of PBS. 50μl of neuraminidase (from Vibrio Cholera, Merk) at a concentration of 50mU/ml, or 50μl of Heparinase III (from Flavobacterium heparinum, Merck) at a concentration of 1U/ml was diluted in serum free media, added to the appropriate wells, and incubated for 1hr at 37°C. Cells were cooled on ice and washed twice with 200μl of PBS. Green Fluorescent Protein (GFP) expressing, replication incompetent viruses were added to the appropriate wells at a concentration of 5000 viral particles per cell, in 100μl of serum free media, at 4°C, and incubated on ice for 1hr. Serum free media alone was added to uninfected control wells. Cells were washed twice with 200μl of cold PBS, complete media added (DMEM, 10% FCS) and incubated for a further 48hrs at 37°C. Cells were then trypsinised and transferred to a 96 well V-bottom plate, washed twice in 200μl of PBS and fixed in 2% paraformaldehyde containing PBS for 20mins before wash, and resuspension in 200μl of PBS.

Samples were analysed by flow cytometry on Attune NxT (ThermoFisher), voltages were set prior to each experiment, for each cell type, using an uninfected cell population treated identically. Data was analysed using FlowJo v10 (FlowJo, LLC), gating sequentially on singlets, cell population, and GFP positive cells. Levels of transdcution were defined as the percentage of GFP positive cells (% +ve), and/or Total Fluorescence (TF), defined as the percentage of GFP positive cells multiplied by the median fluorescent intensity (MFI) of the GFP positive population. These measures are distinct in that % +ve describes the total proportion of cells infected, and TF describes the total efficiency of transgene delivery.

### Surface Plasmon Resonance

Surface plasmon resonance was performed, in triplicate, as previously described, using recombinant HAdV-D49K protein(35). Approximately 5000 RU of Recombinant Human Desmoglein-2 Fc Chimera Protein (R&D Systems, Catalogue number 947-DM-100) was amine coupled to a CM5 sensor chip at a slow flow-rate of 10 μl/min to ensure uniform distribution on the chip surface.

### Competition Inhibition Assay

Competition inhibition assays of antibody binding to cell surface receptors were performed as previously described(35).

### Generation of recombinant fiber knob proteins

Recombinant fiber knob proteins used in transduction inhibition, antibody blocking, and crystallisation experiments were produced as previous described(24, 35). Briefly, pQE-30 vectors containing the sequence of the relevant fiber knob protein, spanning from 13 amino acids preceding the TLW motif to the stop codon, were transformed into SG13009 E.coli harbouring the pREP-4 plasmid. 1L of these E.coli were grown to OD0.6 and protein expression induced with a final concentration of 0.5mM IPTG. E.coli were harvested by centrifugation and resuspended in 50ml lysis buffer (50 mM Tris, pH 8.0, 300 mM NaCl, 1% (v/v) NP40, 1 mg/ml Lysozyme, 1 mM β-mercaptoethanol). Sample was then loaded onto a HisTrap FF Crude column and eluted by imidazole. Fractions determined to contain protein of interest were then concentrated to <1ml total volume and purified by size exclusion chromatography using a Superdex 200 10/300 GL Increase column.

### Fiber knob protein crystallisation and structure determination by X-Ray crystallography

HAdV-D49K, HAdV-D49.KO1.K, and HAdV-D30K were crystallised in the same manner as previously described(24, 35). Both HAdV-D49K and HAdV-D49.KO1.K crystals formed in 0.1M MMT, 25% w/v PEG1500, whilst HAdV-D30K crystallised in 0.1M SPG, 25% w/v PEG 1500. All crystals formed in 2-7 days in sitting drop format. Data collection statistics are described in Table 1 and the structures were solved by molecular replacement using PDB 6FJN.

### Calculation of electrostatic surface potentials and pIs

Electrostatic surface potential and isoelectric points were calculated at pH 7.2 using the PDB2PQR Server (V 2.1.1)(45) as previous described(24).

### RMSD calculation, sequence alignment, and imaging of crystal structures

Alignments were performed using the Clustal Omega multiple sequence alignment algorithm and visualised with BioEdit(46, 47). RMSD calculations were performed using the ‘align’ command in PyMOL 2.0, which was also used to visualise protein structures(48).

## Acknowledgements

ATB was funded by a Tenovus Cancer Care PhD studentship (reference PhD2015/L13) to ALP. GM was supported by a Cancer Research UK ECMC centre award (C7838/A25173). JAD is funded by a Cancer Research UK Biotherapeutic Programme award (reference C52915/A29104) to ALP. EM is supported by a Wellcome Trust ISSF Translational Kickstarter Award (reference 517732) to ALP. RMM was supported by the Cardiff University Research Opportunities Placement (CUROP) to ALP. PJR and ALP are funded by HEFCW. The authors acknowledge the Diamond Light Source for beamtime (proposal MX18812) and the staff of beamline I03 for assistance with diffraction data collection. The authors acknowledge Johanne Pentier for technical assistance with SPR data collection, as well as Aaron Wall and Anna Fuller for assistance with FPLC.

